# Layer-specific reorganization of mnemonic representations in primate retrosplenial cortex during learning

**DOI:** 10.1101/2024.06.30.601381

**Authors:** Niranjan A Kambi, Mohsen Afrasiabi, Jessica M Phillips, Shobha C Kenchappa, Duncan Cleveland, Michelle Redinbaugh, Sounak Mohanta, Bekah Wang, Matt Fayyad, Yuri B Saalmann

**Author notes:** Contributed equally (Mohsen Afrasiabi, Jessica M Phillips). Communicating authors (Niranjan A Kambi, Yuri B Saalmann).

## Abstract

Rapid learning of associations between co-occurring stimuli is essential for episodic memory formation. The retrosplenial cortex (RSC) is strongly interconnected with the hippocampus, and in rodents, the RSC has been shown to support spatial navigation and fear conditioning. Although lesion and neuroimaging studies in humans and macaques have further implicated the RSC in episodic memory, it is unclear how memory representations form and evolve in the RSC. Here we show that representations of memorized contexts in primate RSC form within minutes. These initial representations reorganize as the memory matures, with a shift in the weight of neuronal contributions from superficial to deep RSC layers across an hour and increased local connectivity between deep layer neurons. Because RSC superficial and deep layers represent input and output layers respectively, it suggests that hippocampal inputs provide context information to superficial layers during early learning, and this context information consolidates in deep RSC layers.

## INTRODUCTION

Episodic memory is our ability to reconstruct past experiences. It depends on rapid learning of associations or correlations in our environment with just one or few exposures of the to-be-remembered items. The retrosplenial cortex (RSC) in the medial parietal lobe appears vital for episodic memory (Aggleton and O’Mara, 2022) because, first, neuropsychological studies have shown that damage to RSC leads to retrograde and anterograde amnesia (Valenstein et al., 1987; Gainotti et al., 1998). Secondly, early stages of Alzheimer’s disease are characterized by metabolic changes and atrophy in the posterior cingulate regions including RSC (Minoshima et al., 1997; Nestor et al., 2003; Buckner et al., 2005). Thirdly, RSC is bidirectionally connected with other brain regions involved in episodic memory such as the hippocampus, anterior thalamus, entorhinal and parahippocampal regions (Bubb et al., 2017).

There is also a large body of literature in rodents implicating RSC’s role in spatial memory and navigation (Miller et al., 2014; Mitchell et al., 2018). Neural activity in rodent RSC represents not only specific routes, distances and changes in direction of the animal’s movements in a spatial navigation task, but also sensory cues important for navigation and reward locations in the explored environment (Alexander and Nitz, 2015, 2017; Vedder et al., 2017). Although such representations seem like prerequisites for episodic memory, the “how” and “when” of RSC involvement during learning and formation of episodic memory is unclear.

Rodent studies point to RSC involvement during early stages of learning and memory formation. Conditioning studies involving immediate early gene expression in mice have shown RSC activation occurring within the first few minutes to hours of encoding new information (Katche and Medina, 2017; Baumgärtel et al., 2018). Furthermore, electrophysiological recordings from RSC neurons in rats have shown that the number of cells responding to predictive cues in a T-maze task increases over training sessions which lasted across days (Vedder et al., 2017; Miller et al., 2019). In non-human primates (NHP) performing the object-in-place task which incorporates features of spatial and episodic memory, targeted lesion of the RSC has been shown to cause deficits in memory retrieval after 24 hours of encoding both new (acquired after lesion) and old (before lesion) information, suggesting RSC involvement during memory retrieval (Buckley and Mitchell, 2016). However, there have been no electrophysiological studies in RSC probing memory in NHPs to specify mnemonic representations and their temporal evolution.

Here, we characterize the role of RSC in acquiring episodic-like information, including the dynamics of neural representations, in NHPs during learning. To do this, we used a context memory task in which NHPs learnt the association between a visual array of objects (referred to as context) and a saccade to a specific object in the array. NHPs were exposed to four novel contexts each recording session, and one well-learnt old context, during electrophysiological recordings across RSC layers. We found that RSC in NHPs represents mnemonic experience, and the representations across RSC layers reorganize dynamically during learning.

## METHODS

### Experimental Model and Subject Details

We acquired data from two male monkeys (Macaca mulatta: Noam, 8.7 years old and 9.4 kg body weight; Charlie, 5.9 years old and 8.9 kg body weight). Animal daily needs were maintained by experimenters and animal care staff at the Wisconsin National Primate Research Center (WNPRC), where the macaques were housed. Animal health was monitored and maintained by veterinarians at the WNPRC. The University of Wisconsin-Madison Institutional Animal Care and Use Committee approved all procedures, which conformed to the National Institutes of Health Guide for the Care and Use of Laboratory Animals.

### Stimuli for Memory Task

The stimuli for these experiments consisted of three objects and a central fixation spot (1 degree of visual angle), generated using Presentation software (Neurobehavioral Systems, Inc. Berkeley, CA, USA www.neurobs.com). Two of the three objects were individual colored, alphabetical characters taken from Roman, Devanagari or Kannada scripts (1 degrees of visual angle), while the third object was a colored, irregular-shaped polygon (2 degrees of visual angle). We refer to the alphabetical characters as the saccade target (rewarded) and the distractor (unrewarded). We refer to the colored polygon as the landmark. The saccade target and distractor along with the landmark formed a visuospatial collection referred to as a context (McKenzie et al., 2014; Stark et al., 2018) (Fig 1a).

**Fig 1.**
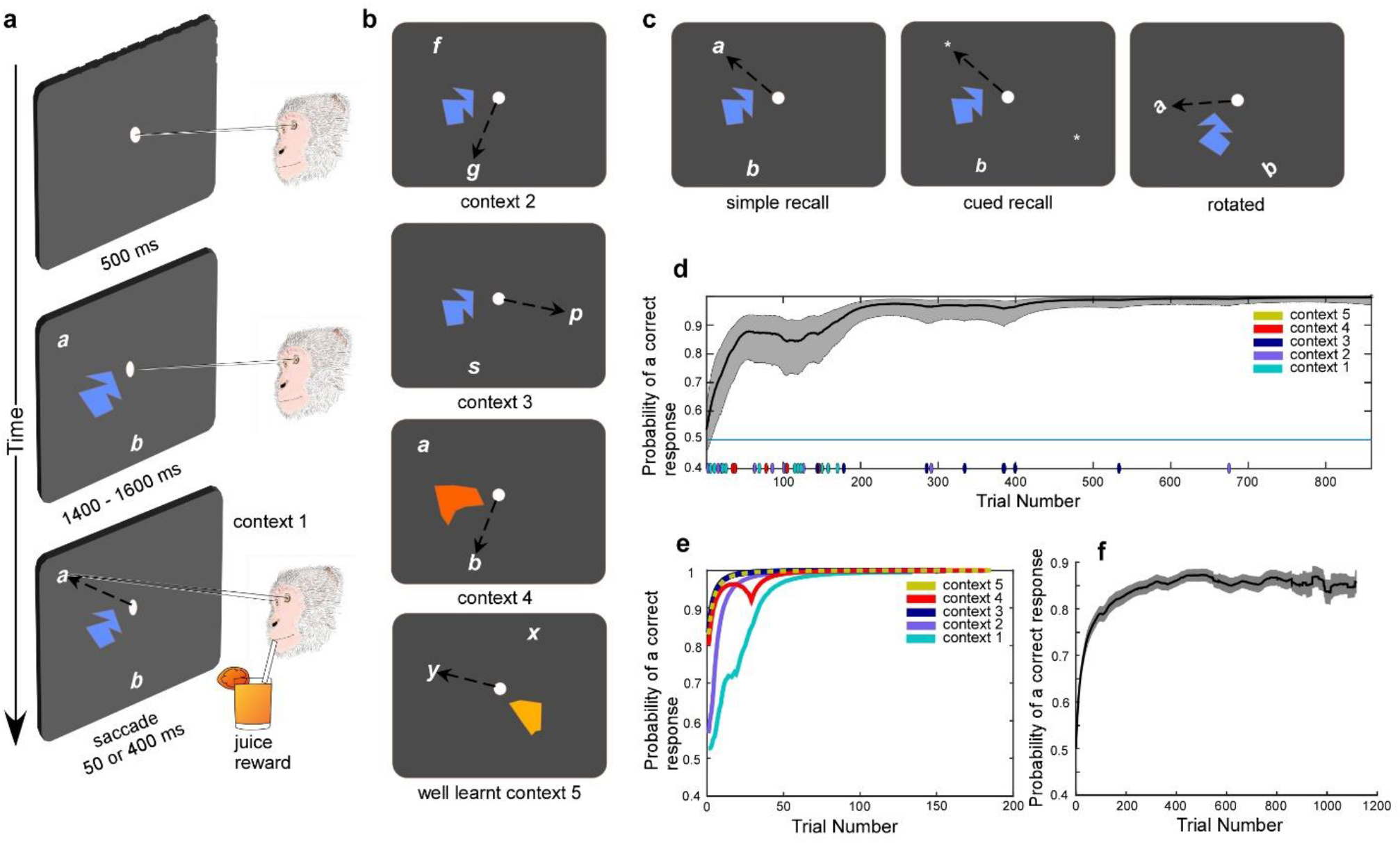
Monkeys learned new contexts and recalled old contexts in the memory task. (a) Trial structure illustrated for context 1. (b) Contexts 2-4 for a particular session; context 5 the same across sessions. (c) Recall trial types. (d) Learning curves for all contexts in typical session. Colored markers show error trials for each context. (e) Learning curve for each context in same session as (d). (f) Average trial-wise learning curve across all sessions.

### Memory Task

We trained macaques to sit in a primate chair (Crist Instruments, Hagerstown, MD, USA www.cristinstrument.com) with their head stabilized to minimize head movements. The task started with a black screen and a central fixation spot. A typical trial started with the subject fixating the central spot for 500 ms after which the context was presented. The context containing the saccade target, distractor, and landmark remained on the screen for a random duration between 1400 ms to 1600 ms while the monkey fixated the central spot. At the end of this duration, the central fixation spot disappeared which was the go cue for the macaque to initiate a saccade to the target, i.e., one of the two characters associated with reward. The macaques learnt by trial and error to saccade to the rewarded target (different auditory feedback indicated correct and incorrect saccades).

The monkeys were presented with four new contexts everyday along with a fifth well-learnt context which was the same everyday, and all contexts were presented in a random, interleaved order. The four new contexts systematically varied from each other in their content objects, to help control for the identity and location of characters/landmarks as well as saccade direction and reward location. Context 1 and 2 have the same landmark and location of characters, but different character identity/color and saccade target, i.e., the rewarded target location in context 2 corresponds to the distractor location in context 1. Context 1 and 3 had the same landmark and distractor location, but different characters and saccade target location, i.e., the rewarded target location was different to that in contexts 1 and 2. Context 4 had the same characters at the same locations as context 1, but different landmark and saccade target, i.e., the rewarded target location in context 4 was the same as that in context 2, but different to that in context 1. Context 5 was completely different in its content of the two characters and the landmark from contexts 1-4, in addition to context 5 being well-learnt (Fig 1b).

Once the animals had performed five consecutive correct trials for a particular context (until this performance was reached, we referred to trials as encoding/training trials), recall versions of the context were presented to test memory (referred to as recall/test trials). Three kinds of recall trials were presented: simple recall, cued recall, and rotated (Fig 1c). The recall trials had the same contexts presented as in the encoding trials. However, in the cued-recall trials, contexts were modified such that the saccade target was replaced by a small placeholder (asterisk character) and an additional placeholder (another asterisk with the same appearance) located at the diametrically opposite location to that of the saccade target. In this case, the monkeys had to saccade to the placeholder at the location of the saccade target by recalling the original contexts. The rotated trials consisted of contexts rotated 45 degrees in the anti-clockwise direction such that the monkeys now had to plan a different saccade to reach the same target as in the encoding trials.

### Behavioral Performance Analysis

To model learning and behavioral performance in our memory task, we applied a state-space paradigm as described in detail in the study by (Smith et al., 2004) and used the MATLAB scripts provided by the authors (https://www.neurostat.mit.edu/behaviorallearning). Briefly, the binary saccadic response in the task was mathematically modeled by a Bernoulli probability distribution while the learning process underlying the performance was modeled by a Gaussian state equation. The probability of making the correct choice in the memory task was termed the learning curve and determined as a function of the trial number using a combination of a forward filter and a fixed smoothing algorithm.

We performed k-means clustering of the probabilities of making correct choice along with the rate (first-order time derivative) and acceleration (second-order time derivative) of these probabilities as additional features for inputs to the clustering algorithm. We used the Silhouette value (Anon, n.d.) as the criterion to get the optimal number of clusters for each session.

### Neuroimaging

We obtained T1-weighted magnetic resonance (MR) images of the monkey brain using the GE MR750 3T scanner (GE Healthcare, Waukesha WI). The imaging was performed under anesthesia which was initially induced using ketamine (up to 20 mg/kg body weight) and atropine sulfate (0.03-0.06 mg/kg) before intubation. General anesthesia was maintained throughout the imaging procedure by administering isoflurane (1%–3% on 1 L/min O2 flow) to the monkeys while their expired carbon dioxide, respiration rate, oxygen saturation, pulse rate and temperature was monitored and recorded.

High-resolution anatomical images of the whole brain were acquired using an inversion-recovery prepared gradient echo sequence with the following parameters: FOV = 128 mm^2^; matrix = 256 × 256; number of slices = 166; voxel dimension = 0.5 mm isotropic; TR = 9.68 ms; TE = 4.192 ms; flip angle = 12°; inversion time (TI) = 450 ms. To generate high quality images, we collected 6-10 T1-weighted images and averaged them using the FSL application (FMRIB Software Library, https://fsl.fmrib.ox.ac.uk/fsl/fslwiki). These scans were used to position the head implant and craniotomy. After implantation, to verify and adjust electrode positions, we collected and averaged 2 T1-weighted images with electrodes *in situ*. We either scanned MRI-compatible linear microelectrode arrays (LMAs, used for recordings; Microprobes Inc https://microprobes.com/) or platinum microelectrodes (later replaced with LMAs inserted along same track; FHC Inc, https://www.fh-co.com/) held in place by customized guide tubes and inserted in the region of interest.

### Implant Surgery

Anesthesia was induced with ketamine (up to 20 mg/kg body weight, i.m.) and maintained with isoflurane (1%–2%) throughout the aseptic procedure. After preparing the dorsal surface of the head for surgery, we made a midline skin incision, freed the skin from the underlying temporalis muscles on both sides, then exposed the dorsal surface of the skull by displacing the temporalis muscles laterally. We drilled eleven 2.5 mm holes in the skull inside the margins of the head implant using a hand-held drill (Thomas recording GmbH, www.thomasrecording.com) and inserted eleven ceramic screws (Thomas recording GmbH) to secure the implant. Dental acrylic (Yates-Motloid Inc, www.yates-motloid.com) was used to build the implant around the screws on the skull. We positioned the recording chamber predominantly on the right hemisphere over brain regions of interest (ROIs, i.e., RSC, hippocampus and anterior thalamus; the hippocampus and anterior thalamus recordings to be part of another study) and the head post on the left hemisphere to hold and stabilize the head. We further secured the implant, including recording chamber and head post, with additional acrylic.

Inside the recording chamber, we drilled three to four 2.5 mm cranial holes in the parietal bone over the ROIs in the right hemisphere. The coordinates of the ROIs were derived from the high-resolution MR structural images acquired before the surgery, using a macaque brain atlas as a reference (Saleem and Logothetis, 2006). Each cranial opening was fitted with a customized plastic guide tube filled with bone wax. We prefabricated the guide tube as a conical frustum tailored to the diameter of the cranial hole and skull thickness (as measured from a model of the skull based on the T1-weighted anatomical images) (Saalmann et al., 2007; Pigarev et al., 2009; Redinbaugh et al., 2020).

### Electrophysiology

We acquired electrophysiological activity from RSC using LMAs (Microprobes, https://microprobes.com/, and Plexon, https://plexon.com/) while monkeys performed the memory task (Fig 2a-c). The LMAs had 24 or 32 channels consisting of Platinum-Iridium electrode contacts (12.5 to 15 μm diameter, with 150 to 200 μm spacing between contacts, and impedance ranging between 0.5-1 MΩ). We recorded wide-band signals from 0.1-7,500 Hz, sampled at 40 kHz, using the OmniPlex data acquisition system with PlexControl software (Plexon). Spike sorting was performed on the filtered signal (250-5,000 Hz) using the Plexon Offline Sorter software (Plexon Inc. https://plexon.com/) for all recording sessions, except one session where we used kilosort 2 software (Pachitariu et al., 2016) with visualization and curation steps in the phy GUI (https://github.com/cortex-lab/phy/). We recorded 307 RSC cells in a total of 49 sessions (22 sessions from monkey N and 29 sessions from monkey C).

**Figure 2.**
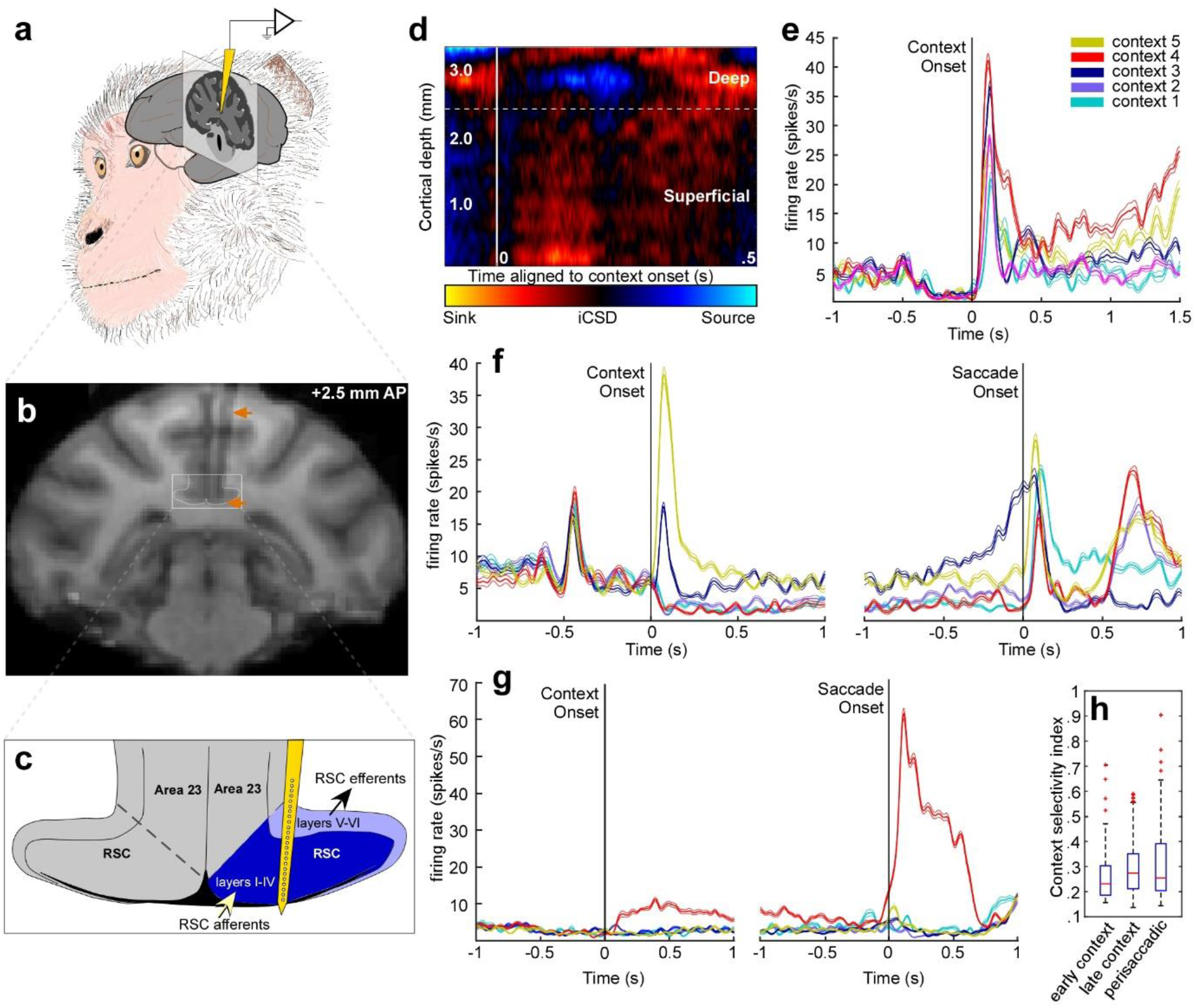
RSC neurons showed context selectivity. (a) Illustration of implanted probe in RSC. (b) Coronal slice of structural MRI with probe (dark shadow, orange arrows) in situ. White rectangle highlights RSC. (c) Schematic showing targeted location of probe contacts (circles on yellow probe) across RSC layers. (d) RSC CSD (color-coded) aligned to context onset in individual session. (e-g) Spike density functions. (e) RSC neuron showing context selectivity. (f) RSC neuron selectively responds to one context at onset and different context during the delay. (g) RSC neuron showing peak, context-selective response in the perisaccadic period. (h) Distribution of context selectivity index for all RSC neurons across three trial epochs.

### Electrode Positioning

We positioned the LMAs in the RSC such that the 4.6 to 4.65 mm long shaft of the probe containing 24 to 32 electrode contacts was nearly perpendicular to the cortical surface. Using the depth measurements from MRI scans with the electrode in the ROI, we aimed for the span of electrode contacts on the probe to maximize the coverage of all layers of RSC. We then adjusted LMA position online to maximize the number of contacts showing single-unit activity. Offline, we used a combination of depth measurements from anatomical scans, single- or multi-unit activity (to help delineate the RSC boundary with white matter), and current source density (CSD) analysis, to divide the electrode contacts into those belonging to superficial versus deep layers in the RSC.

### CSD Analysis and Delineation of RSC Layers

We used inverse CSD analyses to determine the location of electrode contacts and assigned them to the superficial or deep RSC layers (Pettersen et al., 2006). To do this, we used the CSDplotter toolbox for MATLAB (https://github.com/espenhgn/CSDplotter; dt = 1 ms, cortical conductivity value = 0.4 S/m, diameter = 0.5 mm) and calculated the inverse CSD in response to the onset of the context in the memory task. LMAs record LFP, *v*, at N different depths/locations, with spacing *h*, along the z-axis corresponding to a trajectory approximately perpendicular to the RSC cortical surface (Fig 2). The standard CSD, *c*_*st*_, is estimated from the LFPs using the second spatial derivative, i.e.,

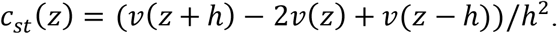

LFPs can also be estimated from given CSDs, represented in matrix form as *V* = ***F**Ĉ*, where *V* is the vector containing the *N* measurements of *v, Ĉ* is the vector containing the estimated CSDs, and ***F*** is an *N* × *N* matrix derived from the electrostatic forward calculation of LFPs from known current sources. The inverse CSD method uses the inverse of ***F*** to estimate the CSD, i.e., *Ĉ* = ***F*** ^−1^***V***. For the step inverse CSD method used here (Pettersen et al., 2006), it is assumed that the CSD is step-wise constant between electrode contacts, so the sources are extended cylindrical boxes with radius *R* and height *h*. In this case, ***F*** is given by:

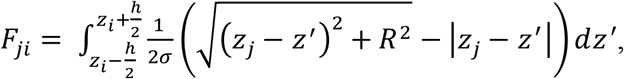

where *σ* is the electrical conductivity tensor, and 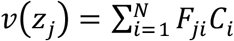 is the potential measured at position *z*_*j*_ at the cylinder center axis due to a cylindrical current box with CSD, *C*_*i*_, around the electrode position *z*_*i*_. The inverse CSD method offers advantages over the standard CSD. The inverse CSD method estimates the CSD around all *N* electrode contacts, whereas the standard CSD method yields estimates around *N* − 2 contacts. Further, the standard CSD requires equidistant contacts, whereas the inverse CSD method does not, which is advantageous when data from a noisy contact may need to be excluded.

We observed an early sink corresponding to spiking in the iCSD plot (Fig 2d) in response to the onset of contexts. We designated the top of this sink as the border between superficial and deep layers. This reflects the fact that the posterior cingulate cortex turns laterally, inverting RSC laminae along the dorso-ventral axis, i.e., the ventral electrode contacts correspond to superficial RSC layers and the dorsal contacts correspond to deep layers (Fig 2b, c). We further validated the layer assignments using the electrode track reconstructions from MR anatomical images as well as the presence or absence of single and multi-unit spiking activity, which also helped mark the boundary between the overlying white matter and underlying grey matter.

Previous studies performing CSD analysis in the RSC (Nixima et al., 2013, 2017; Ferreira-Fernandes et al., 2019; Nitzan et al., 2020) showed an initial sink in superficial layers after stimulation of thalamic/hippocampal afferents or in response to sharp wave ripples in the hippocampus. In line with these studies, our results showed a similar response profile with an early sink after the onset of visual context presentation.

### Univariate Statistical Analyses

#### Context selectivity based on generalized linear models

To determine if neurons are selective for a particular context versus all the other contexts and for other regressors (a landmark or a rewarded target from other landmarks or rewarded targets respectively) we modeled the spike counts of individual neurons as a function of different regressors (contexts, landmarks and rewarded targets) and applied the Generalized Linear Model (GLIM) approach using R software (R Core Team, 2021). The independent/regressor variables were contexts, landmarks, and rewarded targets while the dependent/response variable was spike count during different trial epochs: fixation (0-500 ms before onset of context), early context (25-500 ms post-onset of context), late context (0-900 ms before the cue for saccade), perisaccadic (50 ms prior to and 400 ms after saccade onset) and intertrial interval (ITI; 1000 ms between end of reward/negative feedback of one trial and the start of fixation epoch of the next trial). Since the dependent variable was spike count, we used the Poisson or quasi-Poisson family of distribution in the GLIM depending on whether the distribution of spike counts showed overdispersion. Overdispersion was tested using the *qcc* package in R (Scrucca, 2004).

We used the following regression equation:

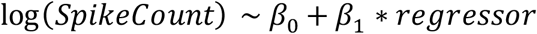

where SpikeCount is the dependent variable and *regressor* refers to regressors: context, landmark, and rewarded targets. *β*_*0*_ is the expected value of spike counts when all regressors are 0. In this paper, the primary focus is context selectivity. *β*_*1*_ is the log of the estimate of the change in spike count associated with change from one context to another, and n refers to the different kinds of contexts presented. The models were run separately for each of the regressors. The p-values from GLIM results were corrected for multiple comparisons using the multcomp package in R (Hothorn et al., 2008). We also determined the chance likelihood that the neurons were selective using the GLIM model by running the above model 1000 times on each neuron’s spike data after random permutation of context labels.

#### Context Selectivity Index

To characterize the neural response to different aspects of the task, we also calculated a selectivity index (SI) using the following formula:

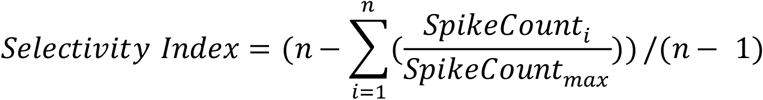

where *n* refers to the total number of contexts (or other aspects of the stimuli), *SpikeCount*_*i*_ is the absolute value of the normalized spike count (subtracted from total spike count during the 500 ms of the central fixation period prior to the onset of the context) and *SpikeCount*_*max*_ is the maximum absolute normalized spike count of the neuron to one of the five contexts. The range of SI was (0,1) where 0 meant the neuron showed no preference for a particular context, while a value approaching 1 meant the neuron preferred one context over all other contexts. We determined the SI for the three major epochs in the trial: early context, late context and perisaccadic. We defined a neuron as context selective if its SI exceeded the 95% quantile of the distribution of 1000 SIs calculated after random shuffling of the labels of contexts (Ku et al., 2021).

#### Firing rate dependence on time

We determined whether the firing rate of single neurons depended on time across a session. We modeled this relationship using the general linear model

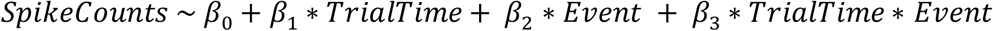

Here, *TrialTime* is a continuous variable referring to the time at which a trial occurred with respect to the total duration of the recording session, and *Event* refers to a discrete variable coding for the epoch in which the spikes were counted with three values corresponding to early context, late context or perisaccadic period.

### Clustering of neural representations

The spike train, i.e, spike count, of a single neuron can be modeled as a generative process with Poisson distribution, with spike timing modeled with an exponential distribution. Though the Poisson process has useful properties, it cannot capture interactions between spikes (self-excitation and self-inhibition) and neurons. For this, we turn to a more general model known as the Hawkes process (Hawkes, 1971). We modeled the spike train of all neurons for a session as a set of time stamps and marks (neuron identification number) for the duration of the task as 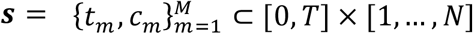 where *T* is the duration of the task, *N* is the number of neurons for a particular recording session, and *M* is the total number of spikes for all neurons in that interval. Sets of discrete events like these are typically modeled as realizations of a marked point process (Daley and Vere-Jones, 2003). Such a process is defined by its nonnegative firing rates, 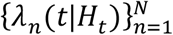, where *H*_*t*_ captures the history of the process through time *t*. For example, the history may include the previous spikes, *H*_*t*_ = {(*t*_*m*_, *c*_*m*_); *t*_*m*_ < *t*}, as well as some external covariates (inputs from other brain areas, which are not considered in this paper). If we consider a small-time window, [*t, t* + Δ*t*) and take the limit as Δ*t* approaches zero, *λ*_*n*_(*t*|*H*_*t*_) × Δ*t* is the expected number of spikes fired by neuron n in the window [*t, t* + Δ*t*). Each neuron has a baseline firing rate, 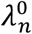. On top of this baseline, each spike from neuron n’ adds a nonnegative impulse response, 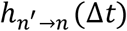, to the subsequent firing rate of the neuron *n*. This allows for spike-driven dynamics that are not possible in Poisson processes. The rate for each neuron is thus given by,

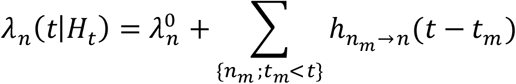

The 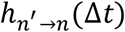 can be decomposed as follows

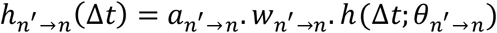

Here, 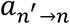 is an entry in the binary adjacency matrix, *A* ∈ {0,1}^*N*×*N*^, and 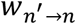 is the corresponding entry in the nonnegative weight matrix, 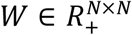. Together these specify the sparsity structure and strength of the interaction network, respectively. The nonnegative function 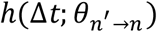 captures the temporal aspect of the interaction. With this formulation, the multi-dimensional Hawkes generative model (MHP) of spikes can be interpreted as a dynamic neural network where *A* is the connectivity matrix resembling the sparse connection of neurons in RSC and *W* can be a proxy for synaptic weights between neurons.

We hypothesized that the network architecture and its underlying parameters would be different in different stages of the experiment due to learning and consolidation of information in RSC. To obtain more insight to the onset of those changes and the sparsity and strengths of the neuronal interaction in each stage, we applied a non-parametric Bayesian clustering approach using a Dirichlet-Hawkes Mixture process for clustering of event streams.

Given a set of spike trains 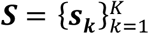, where 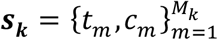 contains a set of neurons *c*_*m*_ ∈ {1,2,. ., *N*} and their timestamps *t*_*m*_ ∈ [0, *T*_*k*_], we modeled the conditional intensity of each sequence of spikes for neuron n belonging to cluster *k* as:

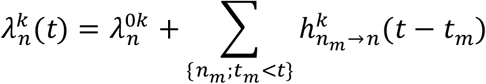

The probability of the appearance of an event sequence ***S*** is:

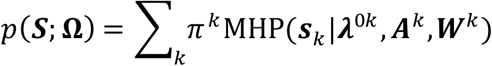

Where *π*^*k*^’s are the probabilities of clusters and MHP(***s***_***k***_ |*λ*^0*k*^, *A*^*k*^, *W*^*k*^) is the conditional probability of spike train *s*_*k*_ given the Hawkes process, which follows the intensity function-based definition in [Daley and Vere-Jones, 2003] and **Ω** is the combination of all parameters. If we denote the latent variables indicating the labels of clusters as matrix ***Z***, we can factorize the joint distribution of all variables and, by using an effective variational Bayesian inference and Expectation-Maximization algorithm (EM), the cluster assignment and Hawkes process’s parameters for each cluster can be derived (Xu and Zha, 2017).

### Multivariate neural decoding

To decode the context from population neural data, we modeled all sessions (n=21) in which we recorded three or more RSC cells. We used spike counts for each 20ms bin as input to the machine learning model. Our machine learning model is designed to capture both spatial (cell interactions) and temporal dependencies for the period of the task aligned to specified events (context onset, saccade onset, etc.). We used a 1D convolutional neural network (CNN) as the first layer to extract rich special features from spike counts of individual neurons and their combined pattern. Since the number of cells recorded in each session is different, we used a max pooling layer after each CNN kernel to make the length of input to the next layer constant. Our kernel sizes were 2, 3, 5, 7, and we used kernels for each size resulting in 100 features as the input to the next layer. To capture the temporal dependency of the firing pattern across the task, we used a recurrent neural network (RNN) with many-to-many architecture to decode the context using enriched features in the previous layer. The proposed RNN consists of a Bidirectional long-short term memory (BiLSTM) layer followed by a multiplicative-attention layer, a fully connected with dropout layer and a softmax layer as output. Considering x(t) as the input at time t, the output of the BiLSTM forward path is calculated as follow:

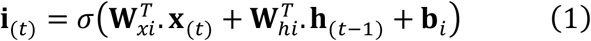

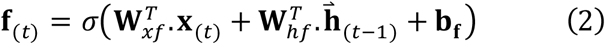

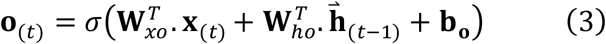

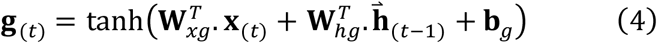

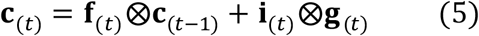

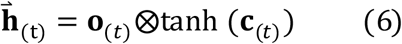

Where **i**_(*t*)_, **f**_(*t*)_, **c**_(*t*)_, **o**_(*t*)_ are the input gate, forget gate, cell gate and output gate respectively, 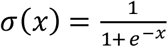. and 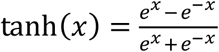, and the output of the backward path is 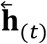. The attention layer output can be calculated as follows:

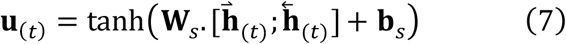

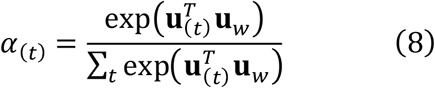

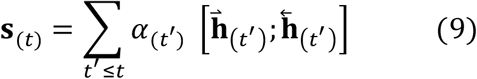

The output of the attention layer was fed to a fully connected layer with dropout followed by a softmax layer which results in class conditional probability as equation (10) specifies.

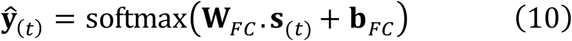

All BiLSTM, attention and FC layers weights and biases were updated through backpropagation in time. Hyperparameters of the model including the number of hidden units for BiLSTM, Attention and FC layers, learning rate, dropout probability, learning rate of stochastic gradient descent, etc., were optimized via a grid search cross validation. The decoding accuracy was calculated with 10-fold cross validation with 60% of trials as the training set and 40% as the test set by shuffling and stratifying to avoid accuracy bias due to imbalanced classes.

To find out how well the model can be generalized in time, we tested the trained model until time *t* with the data at all time points using the stored parameters of the above RNN model. The result of this analysis was used to gauge whether the model trained at time *t* can decode the information at time t’, which gives us insight about how persistent the neural information/representation in the RSC is for the task interval of interest.

## RESULTS

### Context learning within a session and recall across sessions

We used a context memory task which links a particular visual context to a specific action and ensuing juice reward. This is ethologically similar to animals routinely performing actions in specific natural environments to obtain food and other rewards. In our study, two monkeys learned the context memory task (Fig 1A-C) in which they associated a context consisting of a landmark (large shape) and two alphabetic characters (saccade target and distractor) with an eye movement to the saccade target. The task started with encoding trials in which monkeys were presented with 4 novel contexts in random, interleaved sequence and they had to learn by trial and error to saccade to one of the two characters (designated as saccade target) for juice reward. In addition, a well-learnt old context was included in the trial sequence. Each session, the monkeys learnt the four new contexts rapidly, and after they performed five consecutive correct trials for each of the four contexts, recall trials were introduced. The monkeys generalized their initial learning to all three recall trial types – simple, cued and rotated – and we tracked the monkeys’ performance by plotting the probability of a correct response as a function of trial number in a session (Fig 1D-F). This learning curve was derived mathematically by positing an unobservable learning state process which exemplified the dynamics of learning as a function of trial number (Smith et al., 2004). The learning curve showed a sudden increase initially within the first 100 trials after which it stabilized for the rest of the session (typical session, Fig 1D; population, Fig 1F). The monkeys learned each of the four new contexts and maintained the old context, as shown by the learning curve estimates for the five contexts as a function of trial number (Fig 1E), with the best performance early for the well-learnt old context as expected.

### Context selectivity of individual RSC neurons

To probe the neural basis of this context memory, we targeted the RSC (Fig 2A and B) with laminar probes such that the probe contacts spanned RSC layers (Fig 2C). The cytoarchitecture and connectivity of RSC from previous histological studies point to a laminar structure in which the input layers I to IV (from, e.g., hippocampus) occupy almost two thirds of the width of the cortex, with the output layers V and VI (to, e.g., lateral parietal cortex) contained in the remaining one third (see Fig 2C). We used CSD (Fig 2D) to demarcate the border between the input layers (I to IV) and the output layers V and VI, and the presence/absence of spikes and structural brain scans to demarcate the grey matter-white matter boundary.

We recorded a total of 307 cells from both monkeys (n=176 and n=131 in monkeys C and N). To determine the selectivity of cells to individual contexts, we divided the neuronal activity across the entire trial into five epochs: fixation (0-500 ms before onset of context), early context (25-500 ms post-onset of context), late context (0-900 ms before the cue for saccade), perisaccadic (50 ms prior to and 400 ms after saccade onset) and intertrial interval or ITI (1000 ms between end of reward/negative feedback of one trial and the fixation epoch of the next trial). We used two different analytical techniques to show context selectivity, a generalized linear model (GLIM) and context selectivity index (Ku et al., 2021). Based on the GLIM, 34.5% of cells (106 of 307) showed context selectivity, i.e., preference for a single context, in the three epochs starting from the early context presentation to perisaccadic period. Among these cells, 80.2% of cells (85 of 106) showed significantly increased firing to one of the five contexts. Of these 85 cells, 54 (63.5%), exclusively, showed elevated spiking to a specific context, whereas the remaining 31 (36.5%) showed a statistically significant increase in spiking to one context as well as a significant reduction in spiking to another context. The rest of the context selective cells, 31 cells (19.8%) of 106, showed context selectivity by lowering their firing.

Of the context selective cells showing elevated firing, 27.1% (23 of 85 cells) were selective during the early context epoch (Fig 2E), while 45.9% (39 of 85 cells) showed context selectivity during the late context epoch (Fig 2F). The majority of the cells that showed selectivity in the late context epoch were newly recruited (71.8% or 28 of 39), while the remaining 28.2% (11 of 39 cells) of these cells were also active during the early context epoch. Further, during the perisaccadic period, we observed a maximum number (51 of 85 cells) of context selective cells, of which 35 new cells (68.6% of 51 cells) were recruited during this epoch (Fig 2G). Responses during the visually-guided saccade task control did not account for this perisaccadic activity.

Although cells like those in Fig 2G only changed their firing rate in response to one context, other cells like those in Fig 2E and 2F showed firing changes to more than one context (even if there was a significantly different response for one of these contexts). Fig 2E and 2F are cell examples for which different contexts corresponded to different firing rates; the highest firing rate either corresponded to the same context throughout its presentation (Fig 2E), or different contexts depending on the trial epoch (Fig 2F).

To further characterize the selectivity observed in RSC neurons, we calculated a context selectivity index (CSI) and observed that 179 unique cells (58.3% of 307 cells) showed significant CSI (> 95% of the 1000 CSI values calculated for 1000 permutations of randomly shuffled context labels) across all epochs. Of these cells, 51 cells (28.5% of 179 cells) showed significant CSI during the early context epoch, 126 cells (70.4% of 179 cells) showed significant CSI during the late context epoch, and 105 cells (58.7% of 179 cells) showed significant CSI during the perisaccadic epoch (i.e., cells could show significant CSI in more than one epoch). A CSI value close to 1 indicates that the cell is selective for a single context, while values near to 0 imply that the cell is not selective for any individual context. The CSI values of the significantly selective cells ranged from 0.18 (25^th^ percentile value for context onset epoch) to 0.39 (75^th^ percentile for perisaccadic event). Therefore, most cells showed changes in firing rate to more than one context. Such mixed selectivity could be exploited at the cell population level for flexible encoding of many contexts.

### RSC population coding of context

We used a neural decoding approach to investigate how context information is represented in the activity patterns of ensembles of RSC neurons across trial epochs. The decoding accuracy was around chance level (20% for 5 contexts) during the fixation onset epoch as expected. The decoding accuracy increased significantly after the onset of the context presentation (Fig 3A; Fig S1) and remained elevated across the entire time the context was displayed (p = 0.0003, effect size = 6.6). However, the peak accuracy for decoding context occurred immediately after saccade onset (Fig 3B; p =0.0001, effect size = 13.2). Confusion matrices demonstrated the decoding of the five contexts (Fig S2). Therefore, the spiking activity of ensembles of RSC neurons contained information to discriminate between different contexts, during context presentation, in the perisaccadic period, and when receiving rewards without the context on display.

**Figure 3.**
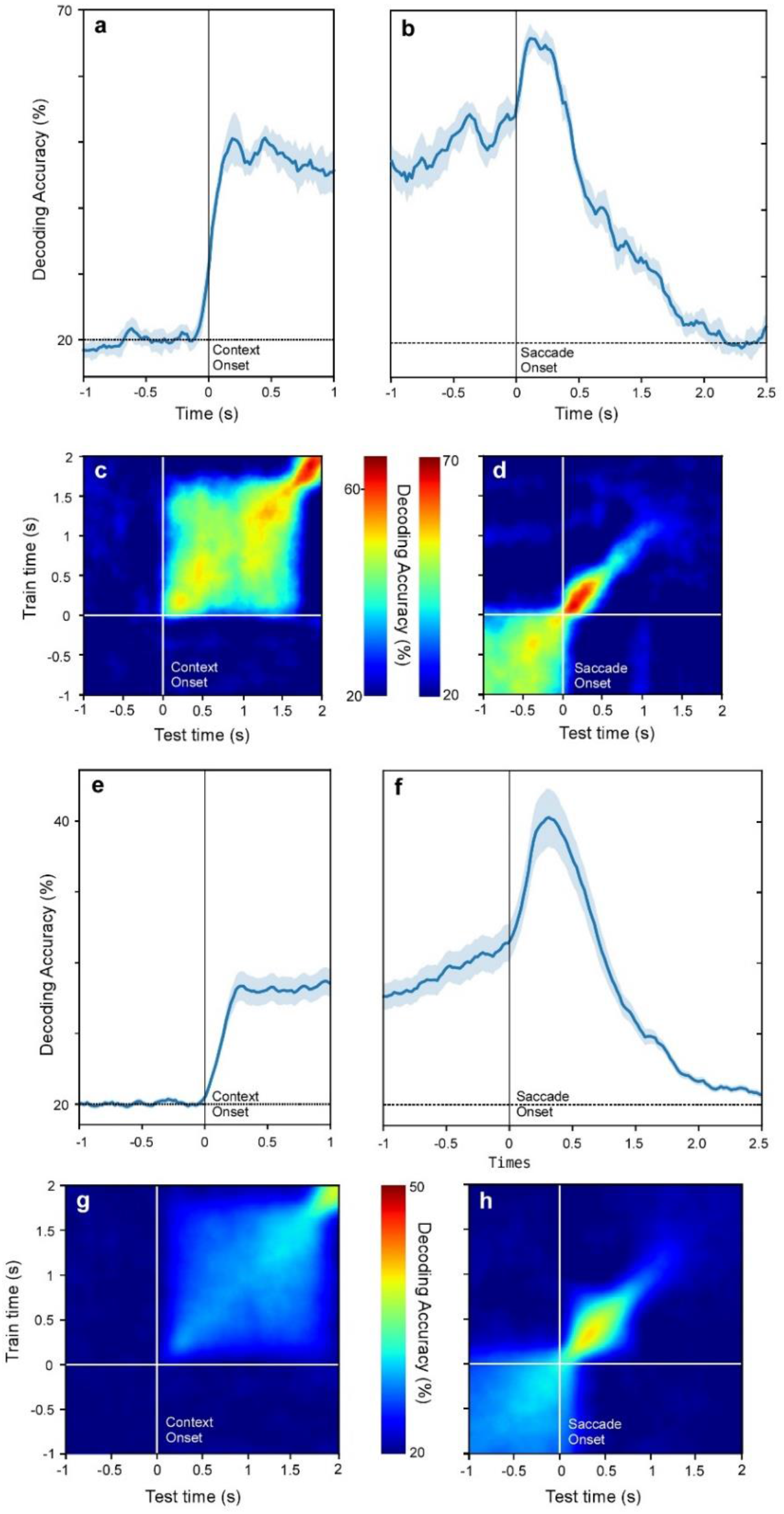
RSC neural ensembles represented context during stimulus presentation, action and reward. (a, b) Decoding accuracy for the five contexts in an individual session. (a) Accuracy increased above chance after onset of context. (b) Decoding accuracy peaked immediately after saccade. (c) Temporal cross training (TCT) plot aligned to context onset for individual session in a. Diagonal represents training and testing at the same time. Note off-diagonal decoding consistent with relatively stable representation during context presentation. (d) TCT plot aligned to saccade onset for individual session in b. Peak post-saccadic performance of the classifier when the training and testing times matched, suggesting more dynamic representation. (e, f) Decoding accuracy for the five contexts aligned to (e) context onset and (f) saccade onset across all sessions. (g) TCT plot aligned to context onset for all sessions, showing off-diagonal decoding. (h) TCT plot aligned to saccade for all sessions, showing highest decoding immediately after saccade when the classifier was trained and tested at same time points.

### RSC population coding dynamics

Next, we investigated the dynamic nature of the population code in RSC, using a temporal cross training (TCT) approach. We tested how well the model trained with context information at any given time during the trial could be used to decode or predict the context at other times during the trial (Fig 3C and D). There was significantly above chance decoding performance in almost all the training/testing pairs starting immediately after the context onset through to the saccade onset (colored square patch in Fig 3C), implying cross-temporal generalization across the entire period. However, the population activity pattern became more dynamic immediately after the onset of the saccade, when the decoding performance was maximal for training and testing during the same time window (colored diagonal patch in Fig 3D). This may have been due to the disappearance of the context from the screen immediately after the saccade, when the context would have to be internally maintained.

We found similar results when averaging the decoding performance across all sessions. Decoding accuracy increased after the context presentation onset (Fig 3E) and reached a maximum after saccade onset (Fig 3F). There was also evidence for cross-temporal generalization across the period from context onset to saccade onset (Fig 3G), but not after saccade onset, when the decoding performance was maximal for training and testing during the same time window Fig 3H). This suggests a relatively stable representation of context until saccade onset, after which context is represented more dynamically.

### Dynamics of firing in individual RSC neurons during context learning

The monkeys learnt four new contexts (contexts 1-4) within an individual session along with recollecting another well-learnt context (context 5). Previously, using c-fos-based optogenetic tagging, stimulation of RSC neurons encoding fear-conditioned contexts elicited fearful reactions independently of the hippocampus in mice during initial learning (Cowansage et al., 2014). However, the dynamics of RSC activity carrying context information during learning is not known. To address this in our context memory task, we used a general linear model to capture how the firing of RSC neurons varied across individual learning sessions; this was done for early/late context and perisaccadic epochs compared to initial fixation. 43 cells in total showed a dependence of firing rate across time. Of these cells, 23 increased their firing with learning and 20 reduced their firing. Notably, these neural changes occurred rapidly, within a recording session (Fig 4). Here we only measured linear changes in the firing of individual cells, which likely underestimates learning-related changes (by not capturing other patterns of change). We next investigated changes in population coding potentially reflecting learning-dependent mechanisms in RSC.

**Fig 4.**
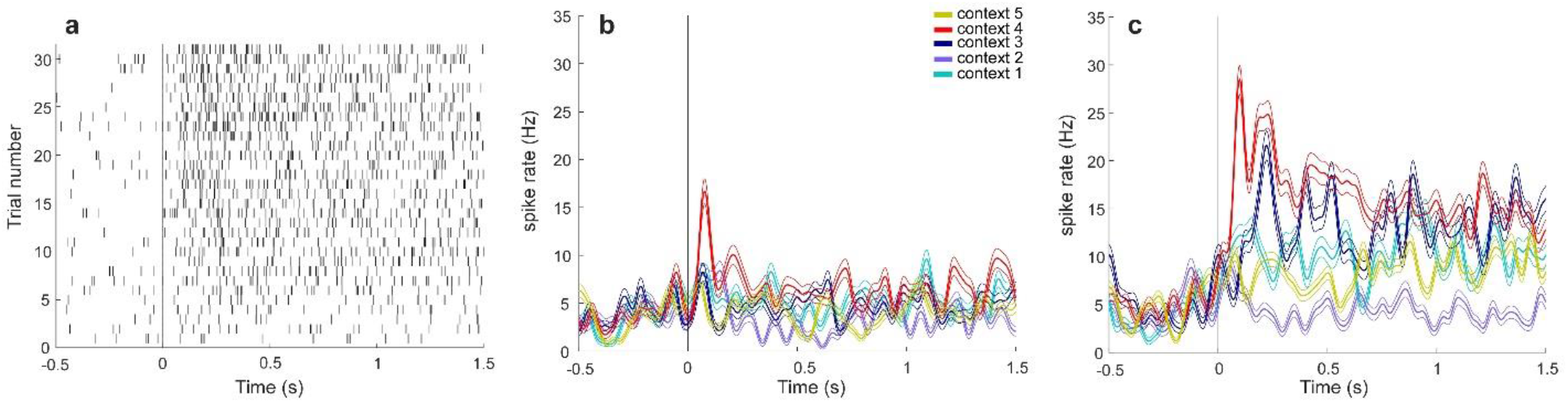
Learning-related activity changes of RSC neurons. (a) Raster plot of RSC neuron showing gradual context-specific increase in firing across trials within a learning session. (b) Spike density functions for all five contexts in early trials. (c) The spike density functions for all five contexts in late trials.

### Correlated timing of neural and behavioral changes

We first characterized how the distributed pattern of firing in RSC cells, representing contexts, developed across time during each of the recording sessions. For this we used a non-parametric Bayesian clustering approach using the Dirichlet-Hawkes Mixture process. The output of this modeling analysis was generally three clusters of trials for each session, organized almost entirely into early, mid, and late trials (Fig 5A).

**Fig. 5.**
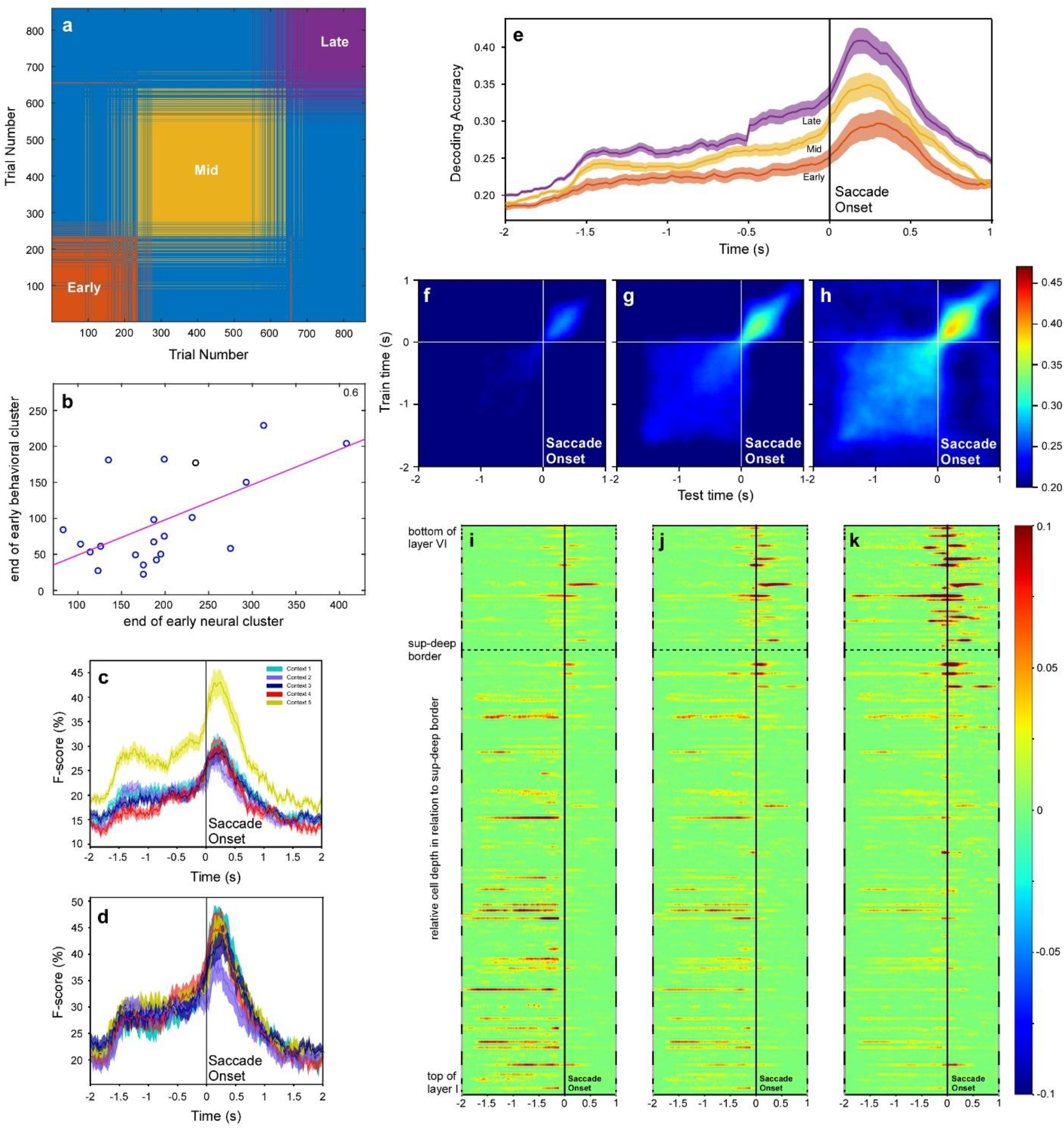
Learning-related changes in context representation across RSC layers. (a) Early, middle and late clusters (orange, yellow and purple, respectively) of RSC neural representations across trials in a typical session. (b) Correlation between early behavioral clusterend and early neural cluster end. Markers represent individual sessions. (c and d) Population average F-score of each context (color-coded) for (c) early and (d) late neural cluster. (e) Population average decoding for early, mid and late neural clusters (color-coded). (f-h) TCT plots for (f) early, (g) mid and (h) late neural clusters. Decoding accuracy color-coded. (i-k) MDA (color-coded) for neurons (rows) organized according to depth in RSC, for (i) early, (j) mid and (k) late clusters.

Next, we tested whether the clustering observed in the neural data correlated with behavior. First, we performed clustering of the behavioral data based on the probability of correct responses, i.e., the learning curve, as well as its first- and second-order time derivatives. The average cluster size across sessions was 2.43: 15 of 21 sessions had two clusters of trials as optimal (as per the Silhouette value criterion), four sessions showed three clusters, and the remaining two sessions showed four and five clusters. Generally speaking, there was an early behavioral cluster(s) followed by a later, more prolonged cluster spanning until the session’s end. This reflects a relatively ‘volatile’, early learning period, followed by a more stable, consolidation period. Next, we calculated the correlation between the end of the early behavioral cluster(s) and the end of the early neural cluster across sessions (Fig 5B). We found a significant correlation (R = 0.6 with p < 0.01 chance of the R occurring at random) indicating a correspondence between learning-related changes in the neural and behavioral data.

### Changes in RSC population coding during context learning

To characterize how the clustering of neural data across a session related to the population representations of context, we performed a decoding analysis for each of the three neural clusters separately and found that the accuracy of classification of contexts increased significantly from early to mid to late clusters, with the late cluster showing maximum decoding accuracy (Fig 5E). This indicates that information regarding context identities in the RSC neural population increased as the behavioral performance improved from early to late trials in a session. To characterize how population representations changed during the learning of specific contexts, we calculated the F-score as a metric for the accuracy of decoding the context. The F-score in early trials was significantly lower for the newly introduced contexts 1-4 compared to the old, well-learnt context 5 (p = 0.00002, effect size = 14.3; Fig. 5C). In contrast, for late trials, there was no significant difference between the new contexts (1-4) and old context (5) (p = 0.004, effect size = 8.1; Fig. 5D). The old, well-learnt context F-score did not differ significantly between the early and late clusters. This suggests that the clustering of the neural data with respect to time (as a proxy of the underlying changes in network interactions) reflects different configurations of the population activity in early, mid and late trial clusters, which correlated with learning to recognize and recall the novel contexts presented to the monkey.

### Changes in RSC population dynamics during context learning

To characterize learning-related changes in the dynamics of neural populations across trial epochs, we performed TCT analyses for the three neural clusters separately (Fig 5 F-H). The decoding accuracy for early, mid and late trials was significantly higher than chance levels for most of the trial duration after the onset of context (p = 0.0005, effect size = 5.8). Further, the difference between early, mid and late decoding accuracies was statistically significant suggesting that the neural representation of context information increased from early to late trials (p = 0.001, effect size = 4.2). For each cluster, we observed peak decoding accuracy immediately after the onset of the saccade (p = 0.003, effect size= 3.5). Decoding accuracy increased from early trials to late trials in the perisaccadic epoch of a trial (p = 0.008, effect size = 2.9). Taken together, this suggests that learning strengthens the population representation of context across all trial epochs from initial sensory stimulation to the recalled action.

### Maturation of memory involves context representations reorganizing across RSC layers

We next tested if the same cells continue to carry context information as the memory matures or if the maturation process transforms the population representation. To do this, we estimated the importance of each cell for decoding the context identities as a proportion of decoding by the entire population of neurons, in each of the early (Fig 5I), middle (Fig 5J) and late (Fig 5K) clusters. Specifically, we calculated the mean decrease in accuracy (MDA) by removing each cell in turn and rerunning the population decoding analysis: The greater the MDA, the greater the importance of that (removed) cell to decoding context.

Firstly, we investigated the dynamics of the information about contexts in the superficial and deep layer neurons. The cells in the superficial layers showed higher MDA during the late context epoch in early trials than that in the mid trials (adjusted p = 1.89263e-10, effect size = 1.206391891) and late trials (adjusted p = 2.93021e-25, effect size = 2.16631256). Further, superficial layer cells had significantly higher MDA during the late context epoch in mid trials when compared to that in the late trials (adjusted p = 7.95267E-09, effect size = 1.092915327). In contrast, deep layer neurons showed increased MDA during the late context epoch as the trials progressed from early to late trials (adjusted p = 0.000132137, effect size = -0.793387137) and similarly from mid to late trials (adjusted p = 0.000398702; effect size = -0.749046011).

Next, we hypothesized that cells in superficial layers of RSC – which receive inputs from other brain areas – carry more information about contexts during the trial (prior to the go cue) than when the animal performs the saccade, for the early trial cluster. On the other hand, for the late trial cluster, when the monkey has learnt the predictive association between the context, action and reward, the information about contexts becomes more crucial around the time the animal performs the action to get reward. In this case, one might expect cells in deep layers to carry more information in the perisaccadic period, as they provide the output from RSC. In line with this hypothesis, superficial layer cells showed higher MDA during the late context epoch as compared to the perisaccadic period in early trials (adjusted p = 9.10313E-27, effect size = 2.030275186). Conversely, in the late trial cluster, the MDA of superficial layer cells decreased in the late context epoch in comparison to that in the perisaccadic period (adjusted p = 3.58972E-27, effect size = - 3.981757952). In contrast to superficial layer cells, deep layer cells showed a consistent pattern of significantly less MDA during the late context epoch as compared to the perisaccadic period in early trials (adjusted p = 1.14331E-09; effect size = -10.69862249), mid (adjusted p = 4.76894E-10; effect size = -4.581673333) and late trials (adjusted p = 9.61445E-13; effect size = -4.560117958).

We directly compared the dynamics of the MDA in the superficial versus deep layers, during the two epochs of interest in early, mid and late trial clusters. During the late context epoch, the neurons in superficial layers showed significantly higher MDA in early (adjusted p = 4.62176E-23; effect size = 1.950037865) and mid trial (adjusted p = 3.73725E-06; effect size = 0.911260752) clusters than the cells in deep layers of RSC. However, in the late trial cluster, the superficial layer cells showed lower MDA (adjusted p = 0.001212904; effect size = -0.715487919) than the deep layer cells during the late context epoch. For the perisaccadic period, the deep layer cells showed a consistently higher MDA than the superficial layer neurons across all the three trial clusters: early (adjusted p = 1.44264E-05; effect size = -5.84903787), mid (adjusted p = 8.64178E-07; effect size = -8.195781151), and late (adjusted p = 9.43965E-08; effect size = -10.30843226). In summary, the MDA increased in the deep layers with the learning of contexts and the maturation of the context memories. This increase in MDA occurred during the delayed context and perisaccadic epochs. These results suggest that the population representation of contexts shifted from the superficial layers of RSC in the early trials to the deep layers in the mid/late trials as the memory matures.

### Changes in RSC network connectivity during context learning

We used a Hawkes process to model network connectivity for the ensemble of RSC cells in each recording session. Connection strength from one cell to another cell can vary from “-1” (inhibitory) to “+1” (excitatory), where “0” is no effect. The connection strength was calculated in 20 ms windows, across the 4 s period spanning 2 s before to 2 s after saccade onset. To calculate graph theoretic metrics, we used the absolute value of the connection strength (ignoring whether inhibitory or excitatory) yielding connection weights from 0 to 1 (i.e., connection weights were not thresholded/binarized). Global efficiency of the whole network increased significantly across a recording session (early vs mid, ANOVA, adjusted p =1.376e-03, effect size = -1.06; early vs late, ANOVA, adjusted p = 1.863e-05, effect size = -1.49; Fig. 6A), which indicates that the efficiency of information transfer across the RSC network increased with learning.

**Figure 6.**
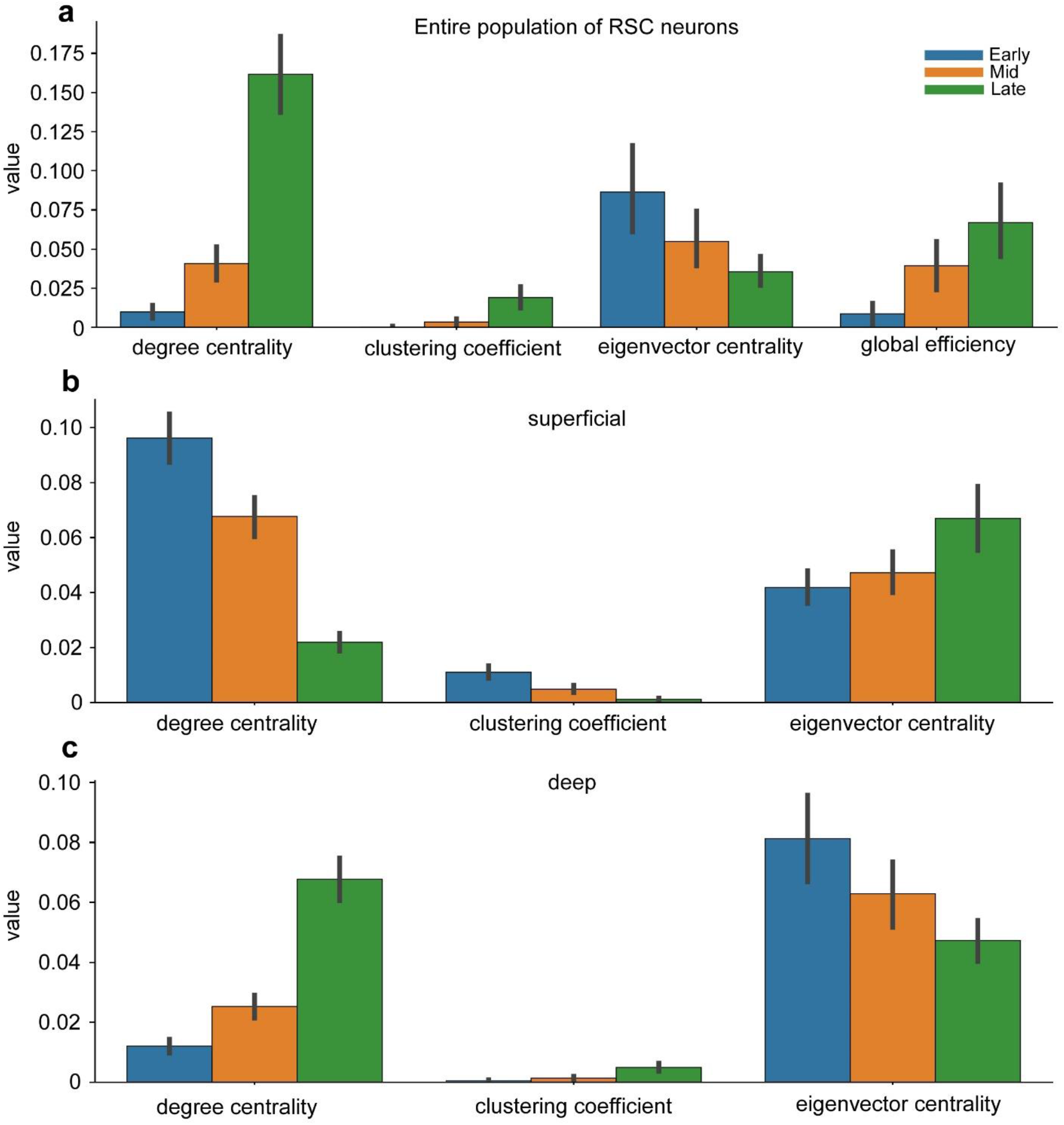
RSC network connectivity for superficial and deep layer neurons changed during context learning. Graph theoretic results for early, mid and later neural clusters, for the (a) entire RSC population, reflecting all cells across all sessions, (b) superficial layer cells and (c) deep layer cells.

Network connectivity was modeled for superficial and deep layer cells all together. To obtain layer-specific graph theoretic measures, we averaged metrics for superficial layer RSC cells and separately averaged metrics for deep layer RSC neurons. For the superficial layers (Fig. 6B), (i) RSC neurons made fewer connections with learning (decreased degree centrality; early vs mid, ANOVA, adjusted p =4.311e-19, effect size = 1.06; early vs late, ANOVA, adjusted p = 3.673e-30; effect size = 2.05; early vs late, ANOVA, adjusted p = 1.179e-14, effect size = 1.39), (ii) there was reduced density of local connections (decreased clustering coefficient; early vs mid, ANOVA, adjusted p = 1.032e-03, effect size = 1.97; early vs late, ANOVA, adjusted p = 6.715e-04, effect size = 2.1; mid vs late, ANOVA, adjusted p = 2.228e-02, effect size = 0.9), but (iii) there was an increase in the influence of superficial RSC neurons on the broader network (increased eigenvector centrality; early vs mid, ANOVA, adjusted p = 2.477e-03, effect size = 1.01; early vs late, ANOVA, adjusted p = 1.626e-07, effect size = 1.85; mid vs late, ANOVA. adjusted p = 1.072e-05, effect size = 1.65). These learning-related changes might reflect more sparse context representations.

For the deep layers (Fig. 6C), (i) RSC neurons made more connections with learning (increased degree centrality; early vs mid, ANOVA, adjusted p = 1.915e-18, effect size = -1.54; early vs late, ANOVA, adjusted p = 5.675e-37, effect size = -2.23; mid vs late, ANOVA, adjusted p = 2.474e-19, effect size = -1.47), (ii) these connections were more local (increased clustering coefficient; early vs mid, ANOVA, adjusted p = 8,461e-03, effect size = -1.73; early vs late, ANOVA, adjusted p = 1.887e-03, effect size = -1.91; mid vs late, ANOVA, adjusted p = 4.387e-02, effect size = -1.04), and (iii) this caused the RSC neurons to become more isolated from the rest of the broader network (decreased eigenvector centrality; early vs mid – ANOVA, adjusted p = 2.320e-03, effect size = -1.17; early vs late, ANOVA, adjusted p = 1.908e-06, effect size = -1.88; mid vs late, ANOVA, adjusted p = 9.793e-06, effect size = -1.43). These changes might serve to make a stored context representation more robust and less prone to interference.

## Discussion

Our results show that context-specific representations in RSC form quickly with learning, within minutes of first exposure. These initial representations reorganize as the memory matures, with the weight of neuronal contributions shifting from superficial layers early to deep layers of RSC across an hour. This reorganization includes deep layer neurons increasing their local connectivity with learning, for a more robust representation of the memorized context. Because the RSC superficial and deep layers represent the input and output layers respectively, it suggests that hippocampal and other inputs give rise to context information in the superficial layers during early learning, and this context information consolidates in the deep layers to enable influence on other brain areas commensurate with learning.

Immediate early gene activation studies in rodents have reported RSC involvement during early learning: RSC activations occurred within minutes after conditioning (Katche and Medina, 2017; Baumgärtel et al., 2018) or changes in the spatial schema to which animals were exposed (Tse et al., 2011). Further, optogenetic activation of cells in RSC which were active during initial fear conditioning led to freezing, pointing to RSC being a cortical storage site for memories (Cowansage et al., 2014). Our study extends these findings in rodents by revealing the neural dynamics of mnemonic representations in RSC of NHPs during the learning of new contexts, including the shift in memory representations from RSC superficial to deep layers. Previous work on layer-specific functions of RSC used calcium imaging in mice moving on a linear treadmill track (Mao et al., 2017). They found that neurons in the superficial layers of RSC sparsely coded for locations in the path similar to place cells in CA1. The place-cell-like responses of the superficial layer RSC neurons were modulated by the presence of tactile stimuli in its path on the treadmill, stabilizing the place-cell-like activity. In our study, we found that superficial layer neurons in RSC contributed most to the decoding of context when NHPs viewed the context before their behavioral report (saccade), but only when first being exposed to new contexts in the early trials of a learning session. After repeated exposures in the mid and late trials, this representation of context information during context viewing slowly diminished, while it strengthened around the time of the behavioral report, particularly in deep layer RSC neurons. A previous study measuring immediate early gene activation in rats found greater sensitivity to spatial and behavioral context respectively in superficial and deep layers of parietal cortex (Burke et al., 2005). Taken together, it may suggest differential mnemonic roles for superficial and deep layers of cortex more broadly, with various linked sensory and spatial aspects of the context initially stored in superficial layers, and behaviorally relevant information later stored in deep layers.

Maturation of engrams in medial frontal cortex of mice suggests increased local and longer-range connectivity between engram cells 28, but not 7, days after contextual fear conditioning (CFC) (Lee et al., 2023). In comparison, we showed increased connectivity in deep layers of RSC, measured as increased degree centrality and clustering coefficient – consistent with stored information becoming more resilient to interference with learning – on the order of 10’s of minutes. This may be due to the repeated presentation of contexts in our task, leading to more rapid changes, or it could reflect RSC’s position as a major output of the hippocampus. Although there were synaptic changes in the medial frontal cortex of mice after 28 days, this was not the case in their RSC, even though information was routed from hippocampus to medial frontal cortex via RSC. In other mouse work, RSC neurons have been reported to be recruited within a day after CFC (Cowansage et al., 2014; de Sousa et al., 2019). This might suggest that RSC plays an intermediate role in systems consolidation, perhaps important for training cortical networks to represent new memories. That said, our well-learnt context was represented in RSC for many weeks across recording sessions. While this may be at least partly due to repeated presentation of this context each session, it does suggest a longer-term role for RSC.

Even though the RSC has been suggested in a number of studies to support spatial navigation and memory in rodents, as well as spatial and episodic memory in humans based on clinical lesions, functional MRI and intracranial EEG recordings (Maguire, 2001; Vann et al., 2009), the relatively few studies of non-human primate RSC have concentrated on sensorimotor functions and seemingly combined data from the RSC with the neighboring and larger posterior cingulate cortex (PCC). These studies have found PCC (and possibly adjoining RSC) responding in the perisaccadic time window – similar to the timing of perisaccadic activity in our study – during visually-guided saccade tasks and the detection of visual events (Olson et al., 1996). Because this activity occurred after saccade onset, it suggests a role in monitoring performance rather than saccade generation (Li et al., 2019). Another study probing PCC (and possibly RSC neurons too) showed subpopulations of cells with saccade target, perisaccadic and reward activity in egocentric or allocentric reference frames for visually-guided saccades (Dean and Platt, 2006), potentially enabling transformations between hippocampal allocentric and cortical egocentric representations.

However, direct comparison of PCC and RSC processing has reported differential processing of self-motion signals, suggesting different roles in, e.g., navigating a context. Compared with strong PCC responses, RSC responded moderately to vestibular perturbation of monkeys as they were linearly translated and rotated passively (Liu et al., 2021). Although our study focused on how whole contexts were represented in the RSC, RSC neurons showed mixed selectivity for the background object (analogous to a landmark to inform saccade direction) in the context, saccade target, direction of saccade, and the onset of events in trials. This is consistent with evidence showing that the RSC responds to the conjunction of sensory, motor, spatial and temporal aspects of tasks (Bucci and MacLeod, 2007; Keene and Bucci, 2008; Robinson et al., 2011; Smith et al., 2012; Fournier et al., 2019), important for representing and recalling context memories, which is impaired when RSC is lesioned (Buckley and Mitchell, 2016). Based on our study, this depends on RSC spiking activity which, at the individual neuron and population levels, represents contexts in superficial layers as soon as context learning starts. As learning progresses, the representations of contexts shifts to deep layer RSC neurons, increasing their local connectivity for more robust representation of the memorized context.

## References

Aggleton JP, O’Mara SM (2022) The anterior thalamic nuclei: core components of a tripartite episodic memory system. Nat Rev Neurosci 23:505–516.

Alexander AS, Nitz DA (2015) Retrosplenial cortex maps the conjunction of internal and external spaces. Nat Neurosci 18:1143–1151.

Alexander AS, Nitz DA (2017) Spatially Periodic Activation Patterns of Retrosplenial Cortex Encode Route Sub-spaces and Distance Traveled. Current Biology 27:1551–1560.e4.

Anon (n.d.) Silhouette criterion clustering evaluation object - MATLAB - MathWorks India. Available at: https://in.mathworks.com/help/stats/clustering.evaluation.silhouetteevaluation.html [Accessed September 5, 2022].

Baumgärtel K, Green A, Hornberger D, Lapira J, Rex C, Wheeler DG, Peters M (2018) PDE4D regulates Spine Plasticity and Memory in the Retrosplenial Cortex. Sci Rep 8:3895.

Bubb EJ, Kinnavane L, Aggleton JP (2017) Hippocampal–diencephalic–cingulate networks for memory and emotion: An anatomical guide. Brain Neurosci Adv 1:2398212817723443.

Buckley MJ, Mitchell AS (2016) Retrosplenial Cortical Contributions to Anterograde and Retrograde Memory in the Monkey. Cerebral Cortex 26:2905–2918.

Buckner RL, Snyder AZ, Shannon BJ, LaRossa G, Sachs R, Fotenos AF, Sheline YI, Klunk WE, Mathis CA, Morris JC, Mintun MA (2005) Molecular, Structural, and Functional Characterization of Alzheimer’s Disease: Evidence for a Relationship between Default Activity, Amyloid, and Memory. J Neurosci 25:7709–7717.

Cooper BG, Mizumori SJY (1999) Retrosplenial cortex inactivation selectively impairs navigation in darkness. NeuroReport 10:625–630.

Cowansage KK, Shuman T, Dillingham BC, Chang A, Golshani P, Mayford M (2014) Direct Reactivation of a Coherent Neocortical Memory of Context. Neuron 84:432–441.

Daley DJ, Vere-Jones D (2003) An introduction to the theory of point processes, 2nd ed. New York: Springer.

Dean HL, Platt ML (2006) Allocentric Spatial Referencing of Neuronal Activity in Macaque Posterior Cingulate Cortex. J Neurosci 26:1117–1127.

Eichenbaum H, Cohen NJ (2014) Can We Reconcile the Declarative Memory and Spatial Navigation Views on Hippocampal Function? Neuron 83:764–770.

Ferreira-Fernandes E, Pinto-Correia B, Quintino C, Remondes M (2019) A Gradient of Hippocampal Inputs to the Medial Mesocortex. Cell Reports 29:3266–3279.e3.

Fournier DI, Monasch RR, Bucci DJ, Todd TP (2020) Retrosplenial cortex damage impairs unimodal sensory preconditioning. Behavioral Neuroscience 134:198–207.

Gainotti G, Almonti S, Di Betta AM, Silveri MC (1998) Retrograde amnesia in a patient with retrosplenial tumour. Neurocase 4:519–526.

Hawkes AG (1971) Spectra of some self-exciting and mutually exciting point processes. Biometrika 58:83–90.

Hothorn T, Bretz F, Westfall P (2008) Simultaneous Inference in General Parametric Models. Biometrical Journal 50:346–363.

Katche C, Medina JH (2017) Requirement of an Early Activation of BDNF/c-Fos Cascade in the Retrosplenial Cortex for the Persistence of a Long-Lasting Aversive Memory. Cerebral Cortex 27:1060–1067.

Ku S, Hargreaves EL, Wirth S, Suzuki WA (2021) The contributions of entorhinal cortex and hippocampus to error driven learning. Commun Biol 4:1–12.

Liu B, Tian Q, Gu Y (2021) Robust vestibular self-motion signals in macaque posterior cingulate region. eLife 10:e64569.

Maguire EA (2001) The retrosplenial contribution to human navigation: a review of lesion and neuroimaging findings. Scand J Psychol 42:225–238.

Mao D, Kandler S, McNaughton BL, Bonin V (2017) Sparse orthogonal population representation of spatial context in the retrosplenial cortex. Nat Commun 8:243.

McKenzie S, Frank AJ, Kinsky NR, Porter B, Rivière PD, Eichenbaum H (2014) Hippocampal Representation of Related and Opposing Memories Develop within Distinct, Hierarchically Organized Neural Schemas. Neuron 83:202–215.

Miller AMP, Mau W, Smith DM (2019) Retrosplenial Cortical Representations of Space and Future Goal Locations Develop with Learning. Current Biology 29:2083–2090.e4.

Miller AMP, Vedder LC, Law LM, Smith DM (2014) Cues, context, and long-term memory: the role of the retrosplenial cortex in spatial cognition. Front Hum Neurosci 8:586.

Minoshima S, Giordani B, Berent S, Frey KA, Foster NL, Kuhl DE (1997) Metabolic reduction in the posterior cingulate cortex in very early Alzheimer’s disease. Ann Neurol 42:85–94.

Mitchell AS, Czajkowski R, Zhang N, Jeffery K, Nelson AJD (2018) Retrosplenial cortex and its role in spatial cognition. Brain Neurosci Adv 2:2398212818757098.

Nestor PJ, Fryer TD, Ikeda M, Hodges JR (2003) Retrosplenial cortex (BA 29/30) hypometabolism in mild cognitive impairment (prodromal Alzheimer’s disease). Eur J Neurosci 18:2663–2667.

Nitzan N, McKenzie S, Beed P, English DF, Oldani S, Tukker JJ, Buzsáki G, Schmitz D (2020) Propagation of hippocampal ripples to the neocortex by way of a subiculum-retrosplenial pathway. Nat Commun 11:1947.

Nixima K, Okanoya K, Ichinohe N, Kurotani T (2017) Fast voltage-sensitive dye imaging of excitatory and inhibitory synaptic transmission in the rat granular retrosplenial cortex. Journal of Neurophysiology 118:1784–1799.

Nixima K, Okanoya K, Kurotani T (2013) Current source-density analysis of intracortical circuit in the granular retrosplenial cortex of rats: A possible role in stimulus time buffering. Neuroscience Research 76:52–57.

Olson CR, Musil SY, Goldberg ME (1996) Single neurons in posterior cingulate cortex of behaving macaque: eye movement signals. Journal of Neurophysiology 76:3285–3300.

Osawa A, Maeshima S, Kunishio K (2008) Topographic Disorientation and Amnesia due to Cerebral Hemorrhage in the Left Retrosplenial Region. Eur Neurol 59:79–82.

Pachitariu M, Steinmetz NA, Kadir SN, Carandini M, Harris KD (2016) Fast and accurate spike sorting of high-channel count probes with KiloSort. In: Advances in Neural Information Processing Systems. Curran Associates, Inc. Available at: https://proceedings.neurips.cc/paper_files/paper/2016/hash/1145a30ff80745b56fb0cecf65305017-Abstract.html [Accessed November 7, 2023].

Pettersen KH, Devor A, Ulbert I, Dale AM, Einevoll GT (2006) Current-source density estimation based on inversion of electrostatic forward solution: Effects of finite extent of neuronal activity and conductivity discontinuities. Journal of Neuroscience Methods 154:116–133.

Pigarev IN, Saalmann YB, Vidyasagar TR (2009) A minimally invasive and reversible system for chronic recordings from multiple brain sites in macaque monkeys. Journal of Neuroscience Methods 181:151–158.

Pothuizen HHJ, Aggleton JP, Vann SD (2008) Do rats with retrosplenial cortex lesions lack direction? European Journal of Neuroscience 28:2486–2498.

R Core Team (2021) R: A language and environment for statistical computing. R Foundation for Statistical Computing, Vienna, Austria. URL. Available at: https://www.R-project.org/.

Redinbaugh MJ, Phillips JM, Kambi NA, Mohanta S, Andryk S, Dooley GL, Afrasiabi M, Raz A, Saalmann YB (2020) Thalamus Modulates Consciousness via Layer-Specific Control of Cortex. Neuron 106:66–75.e12.

Robinson S, Keene CS, Iaccarino HF, Duan D, Bucci DJ (2011) Involvement of retrosplenial cortex in forming associations between multiple sensory stimuli. Behav Neurosci 125:578–587.

Saalmann YB, Pigarev IN, Vidyasagar TR (2007) Neural Mechanisms of Visual Attention: How Top-Down Feedback Highlights Relevant Locations. Science 316:1612–1615.

Saleem K, Logothetis N (2006) A Combined MRI and Histology Atlas of the Rhesus Monkey Brain in Stereotaxic Coordinates - 1st Edition. Available at: https://www.elsevier.com/books/a-combined-mri-and-histology-atlas-of-the-rhesus-monkey-brain-in-stereotaxic-coordinates/saleem/978-0-12-372559-2 [Accessed January 29, 2022].

Scrucca L (2004) qcc: an R package for quality control charting and statistical process control. Available at: https://cran.r-project.org/doc/Rnews/.

Smith AC, Frank LM, Wirth S, Yanike M, Hu D, Kubota Y, Graybiel AM, Suzuki WA, Brown EN (2004) Dynamic Analysis of Learning in Behavioral Experiments. J Neurosci 24:447–461.

Stark SM, Reagh ZM, Yassa MA, Stark CEL (2018) What’s in a Context? Cautions, limitations, and potential paths forward. Neurosci Lett 680:77–87.

Tse D, Takeuchi T, Kakeyama M, Kajii Y, Okuno H, Tohyama C, Bito H, Morris RGM (2011) Schema-dependent gene activation and memory encoding in neocortex. Science 333:891–895.

Valenstein E, Bowers D, Verfaellie M, Heilman KM, Day A, Watson RT (1987) Retrosplenial amnesia. Brain 110 (Pt 6):1631–1646.

Vann SD, Aggleton JP (2004) Testing the importance of the retrosplenial guidance system: effects of different sized retrosplenial cortex lesions on heading direction and spatial working memory. Behavioural Brain Research 155:97–108.

Vann SD, Aggleton JP, Maguire EA (2009) What does the retrosplenial cortex do? Nat Rev Neurosci 10:792–802.

Vedder LC, Miller AMP, Harrison MB, Smith DM (2017) Retrosplenial Cortical Neurons Encode Navigational Cues, Trajectories and Reward Locations During Goal Directed Navigation. Cerebral Cortex 27:3713–3723.

